# Deep Learning Enhanced Hyperspectral Fluorescence Lifetime Imaging

**DOI:** 10.1101/2020.01.06.896092

**Authors:** Marien Ochoa, Alena Rudkouskaya, Ruoyang Yao, Pingkun Yan, Margarida Barroso, Xavier Intes

## Abstract

Acquiring dense high-dimensional optical data in biological applications remains a challenge due to the very low levels of light typically encountered. Single pixel imaging methodologies enable improved detection efficiency in such conditions but are still limited by relatively slow acquisition times. Here, we propose a Deep Learning framework, NetFLICS-CR, which enables fast hyperspectral lifetime imaging for *in vivo* applications at enhanced resolution, acquisition and processing speeds, without the need of experimental training datasets. NetFLICS-CR reconstructs intensity and lifetime images at 128×128 pixels over 16 spectral channels while reducing the current acquisition times from ∼2.5 hours at 50% compression to ∼3 minutes at 99% compression when using a single-pixel Hyperspectral Macroscopic Fluorescence Lifetime Imaging (HMFLI) system. The potential of the technique is demonstrated *in silico, in vitro* and *in vivo* through the monitoring of receptor-ligand interactions in mice liver and bladder and further imaging of intracellular drug delivery of the clinical drug Trastuzumab in live animals bearing HER2-positive breast tumor xenografts.

---

Over the last two decades, computational imaging has experienced an explosive growth thanks to its aptitude to overcome limitations of conventional imaging methods^1^. Especially, optical computational imaging has led to the development of new imaging techniques that have greatly impacted numerous fields including material sciences, computer vision and biomedical sciences^2^. These technical breakthroughs have been stimulated by the wide availability of new optical components such as light modulators and/or sensitive digital sensors. Moreover, the emergence of the theoretical framework of compressive sensing has empowered the implementation of efficient computational imaging systems^3^. Among all technical approaches, structured light techniques have played a significant role in advancing this field^4,5^. For instance, the incorporation of spatial light modulators in the optical chain has led to improved resolution in microscopy^6^, enabled the reconstruction of 4D phase amplitude data^7,8^, facilitated hyperspectral microscopy^9,10^ or enabled fast optical tomography of thick specimens^11,12^. Nevertheless, there is still a strong incentive to develop novel imaging techniques for biomedical applications, that can acquire high-dimensional data (space, time, phase and spectrum), with increased information content, such as required in highly multiplexed molecular imaging.^13^ However, biological applications, especially *in vivo* fluorescence-based imaging applications are characterized by faint signals due to the scattering and absorptive nature of intact (non-cleared) tissues. One computational imaging approach that is well positioned to tackle such low signal conditions is single-pixel imaging^14^. To date, the single pixel imaging approach has been implemented for various applications ranging from astronomy to medical imaging.^15^ A single-pixel camera is typically based on an inverse problem that aims at reconstructing 2D images from measurements that sample the object of interest with a series of spatially encoded patterns (structured light) and associating these known masks with the corresponding 1D intensity measured with a single detector. Such arrangement allows the production of imaging systems with lower cost, improved detection efficiency, lower dark counts and/or fast time-resolved capabilities.

Recently, single-pixel imaging’s potential to concurrently acquire spatial, temporal and spectral data sets at the macroscopic level has been demonstrated via the implementation of single-pixel based Hyperspectral Macroscopic Fluorescence Lifetime Imaging (HMFLI). It integrates an ultra-fast laser, spatial light modulators, and a Time Correlated Single Photon Counting (TCSPC) spectrophotometer.^16^ This imaging platform enables the acquisition of dense hyperspectral time-domain data for functional imaging or lifetime-based molecular imaging for *in vivo* applications, both in 2D or 3D.^17,18^ However, despite the multiple advantages of single-pixel arrangements, one major drawback is the need to acquire large sets of spatially coded measurements to obtain accurate and high resolution image reconstructions. Such requirement potentially leads to long acquisition times or ultra-weak signal conditions as well as long computational times. Moreover, due to the ill-posed nature of the inverse solvers, it is typical to employ advanced regularization techniques, which can be dependent on *post hoc* selection of tuning parameters. In this regard, we have recently proposed the use of deep learning, to simultaneously reconstruct intensity and lifetime single pixel images, without the use of parameter constrained inverse solvers.^19^ However, this architecture is still dependent on inputting a large set of spatially coded measurements to form the images. To overcome this obstacle and accelerate bed-side translation of single-pixel imaging, we propose NetFLICS-CR, a novel deep learning framework that reduces the acquisition times for HMFLI by more than one order of magnitude. Herein, NetFLICS-CR is tested and validated within the framework of HMFLI for preclinical small animal studies. HMFLI can acquire dense spatial, spectral and temporal data cubes for multiplexed fluorescence molecular imaging.^16^ The technique is poised to advance the drug development pipelines by enabling the simultaneous monitoring of multiple biomarkers as well as the quantification of targeted drug cellular delivery via the measurement of near infrared (NIR) labeled receptor-ligand engagement via Föster Resonance Energy Transfer (FRET) HMFLI.^20^ First, we report on the design and *in silico* training as well as validation of NetFLICS-CR. Then, we establish NetFLICS-CR robustness by reconstructing controlled *in vitro* experimental data never used during the *in silico* training phase. Subsequently, NetFLICS-CR is used to quantify iron-bound transferrin (Tf) based target engagement in the liver and bladder of live, intact mice. Lastly, we report its applicability to monitor *in vivo* drug delivery via HMFLI FRET in mice bearing breast cancer tumor xenografts. Specifically, intracellular delivery of trastuzumab (TZM) was quantified in human epidermal growth factor receptor 2 (HER2)-positive tumors in live intact mice. TZM is an antibody that binds to HER2 in surface of cancer cells. Notably, TZM is a FDA-approved drug for the treatment of metastatic breast and gastric cancer that overexpress HER2^21^.

Single pixel imaging retrieves fluorescence time domain (TD) information of the sample plane ***x*** through inverse solving **m*(t) = Px(t)*** (**Eq. 1**) where ***P*** represents the sensitivity matrix (weights) of patterns used for the acquisition of ***m(t)***, composed of an 1D measurement per pattern ***P*** over time **t**, so that the number of 1D measurements needed for reconstruction equals the number of pixels at the desired resolution. This time consuming process can be alleviated by reducing the number of acquisition patterns through compressive sensing (CS) strategies based on specific illumination basis and inverse solvers like the ubiquitously employed TVAL3^22,23^. In order to reconstruct the image plane through this conventional CS workflow, which will be called TVRecon throughout the text, the spatial information is first retrieved and then, the fluorescence Temporal Point Spread Functions (TPSFs) at each pixel are model-fitted to retrieve the lifetime parameters of interest. This process is further explained in **Supplementary Material Section 1**. One important aspect of this workflow is that TVRecon is highly dependent on user defined parameters for TD reconstruction, resulting in an extra optimization step that requires a separate Least-Squares-minimization (LSM) algorithm for lifetime reconstruction. We recently demonstrated this sequential image reconstruction tasks could be performed efficiently and simultaneously using deep learning (DL). Indeed, with NetFLICS^19^, a Convolutional Neural Network (CNN), single pixel raw fluorescence data can be reconstructed into intensity and lifetime images in a single workflow and without the need of parameter optimization. Compared to the TVRecon workflow and when using 50% of the patterns for a 32×32 resolution, NetFLICS produced higher quality reconstructions even at low photon counts and at computational speeds 4 orders of magnitude faster. Despite these promising initial results, NetFLICS only reconstructs 32×32 images at a fixed compression ratio of 50% (512 measurements out of 1024). Even-though the increase in processing speed is essential to apply HMFLI for *in vivo* settings, it is necessary to simultaneously increase resolution beyond 32×32 pixels while decreasing the acquisition time.

Herein, a new CNN architecture, NetFLICS-CR, as displayed in **Figure 1 (a)**, is proposed to retain the fast processing speed for simultaneous intensity and lifetime retrieval but reconstruct 128×128 resolutions while using high compression ratios (CRs) to reduce acquisition time. CRs are defined as the ratio of the number of acquired patterns over the full number of patterns in the compressive basis for a determined resolution. For instance, for a 128×128 resolution, a 90% CR uses 1640 patterned measurements out of 16,384 while a 99% CR uses only 163 out of 16,384. The architecture of NetFLICS-CR is composed of an input segment that convolves along the temporal dimension. The output of this 1D convolution layer is fed to the two main sections of the network: intensity and lifetime branches. The intensity branch is composed of a single ResBlock^24^ and ReconBlock^25^ that yields the reconstructed 128×128 intensity image, with kernel numbers and sizes adjusted to account for the resolution increase. The lifetime branch contains a ResBlock, a double ReconBlock structure and an additional 1D convolutional layer continued by a 2D separable convolution, which help to extract lifetime features across the time dimension. The 1D convolutional layers are followed by batch normalization^26^ and ReLU^27^ activations. For training, a similar strategy as laid in ^18^ is employed. Briefly, the EMNIST dataset is used as the spatial model to generate 128×128 pixels images. Then, at each pixel, fluorescence time-domain data is simulated using a mono-exponential decay model convolved with an HMFLI instrumental response function (IRF). The architecture and training process are detailed in **Supplementary Material Section 1**.

**Fig. 1.**
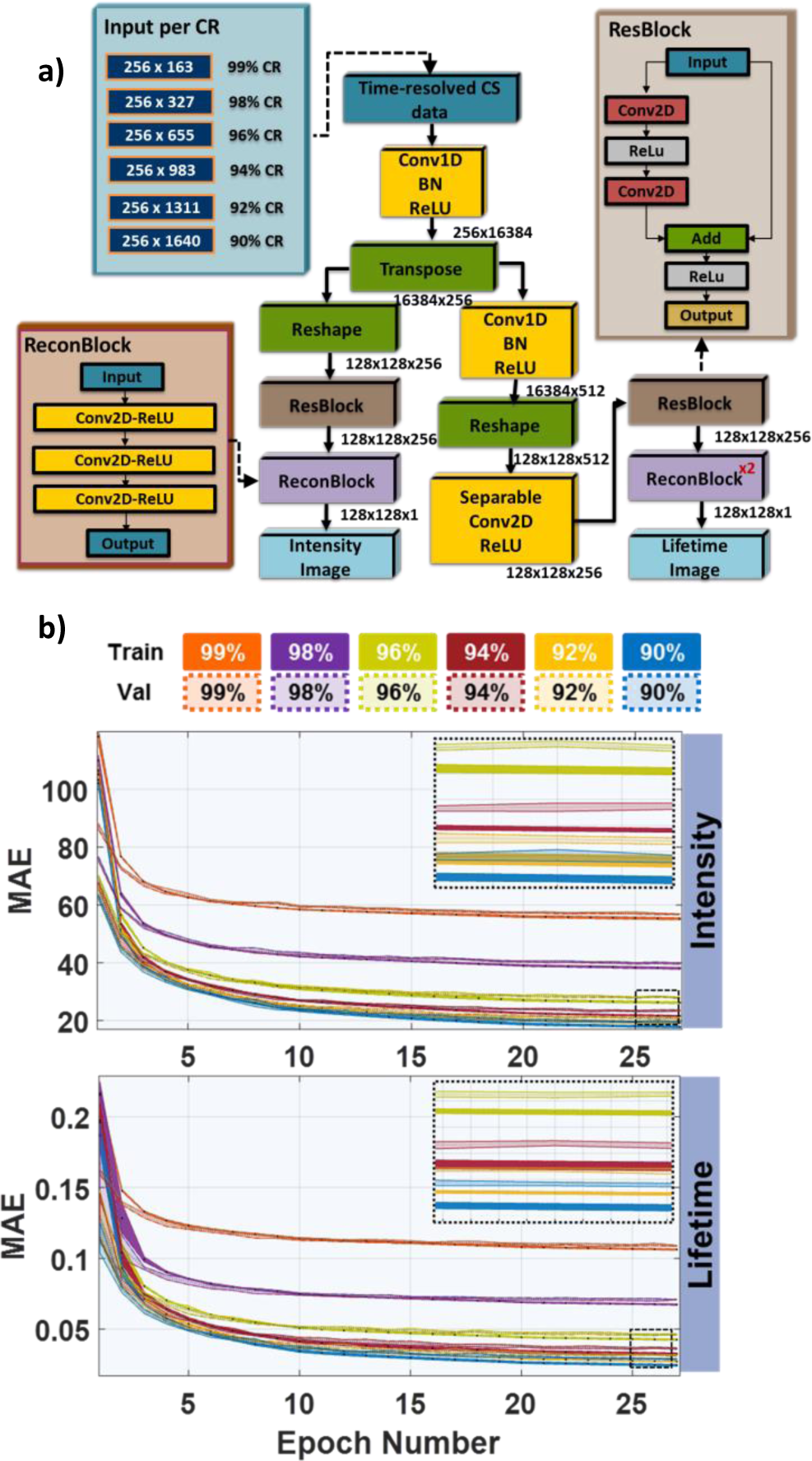
NetFLICS-CR architecture: a) Structure for reconstruction of intensity and lifetime images at different compression ratios. b) Intensity and lifetime training - validation curves at different compression ratios for 5 different training routines.

For network training and validation per CR, 40,000 *in silico* TD samples were generated in batch sizes of 10, with 32,000 used in training and 8,000 in validation (a sample refers to an EMNIST image with an intensity and fluorescence decay range per pixel; therefore, a 128×128×256 array). The single pixel measurements are generated by using **Eq. 1**. To establish the compression of NetFLICS-CR, only a subset of the pattern basis (***P)*** is employed for generating the single pixel measurements. In all cases, the Hadamard Ranked basis (patterns ranked from low to high frequency) ^22^ is used. Therefore, a 99% CR corresponds to only using the first 1% of the ranked Hadamard basis, while 90% CR to using the first 10%. The number of raw single pixel measurements considered per CR is varying as displayed in **Fig. S1**. Note that the CR operates on the spatial dimension. Overall, the number of kernels, their sizes and the main segments of NetFLICS-CR, follow the same scheme for all CRs. The training and validation for both intensity and lifetime branches as evaluated through Mean Absolute Error (MAE) are displayed in **Figure 1(b)** per CR. Additionally, the process was repeated 5 times per CR to evaluate training robustness. The results indicate, for both intensity and lifetime, that as the CR increases the MAE also increases, which is expected as there is less data to reconstruct from. The training and validation curves converge in similar MAE values for the 5 different trainings for each CR, as shown through the solid color shading for training curves and transparent color for validation curves (**Figure 1 (b)**). In order to evaluate NetFLICS-CR accuracy in terms of image reconstruction at each compression ratio, 400 new TD *in silico* samples (different from the training and validation set), were reconstructed. The same samples were reconstructed with the two-step TVRecon procedure, yielding an intensity and lifetime image per CR. The results of both methods were compared versus ground-truth through the Structural Similarity Index Matrix (SSIM) value, which for an ideal reconstruction should equal 1.

The 128×128 intensity and lifetime reconstructions for the three highest compression ratios are displayed for each method in **Figure 2(a)** for an example data set. The average SSIM for the sample sets is shown in **Figure 2(b)** for intensity and lifetime, NetFLICS and TVRecon methods. Since NetFLICS-CR was trained 5 times per compression, one SSMI value per each training and CR is plotted. SSIM values for both intensity and lifetime are higher than 0.8 at 99% CR and the SSIM values increase close to 0.9 as the CR decreases to 90%, which is expected due to an increase in information content. Of note, SSIM values for lifetime are higher than those for intensity. This might be attributed to the higher range in simulated intensity weights compared to lifetimes (also representative to experimental conditions). On the other hand, for all cases, TVRecon SSIM values are below 0.8 at all CRs. Note that the TVRecon method requires selection of the best regularization parameters, which were optimized to values yielding the highest SSIM intensity reconstructions versus ground-truth. After optimization and TD reconstructions, LSM was applied to yield lifetime images. This process is shown and further explained in **Fig. S2(c).** Overall, the *in silico* results with known ground-truth reconstructions indicate a better performance for NetFLICS-CR with SSIM values always above 0.8.

**Fig. 2.**
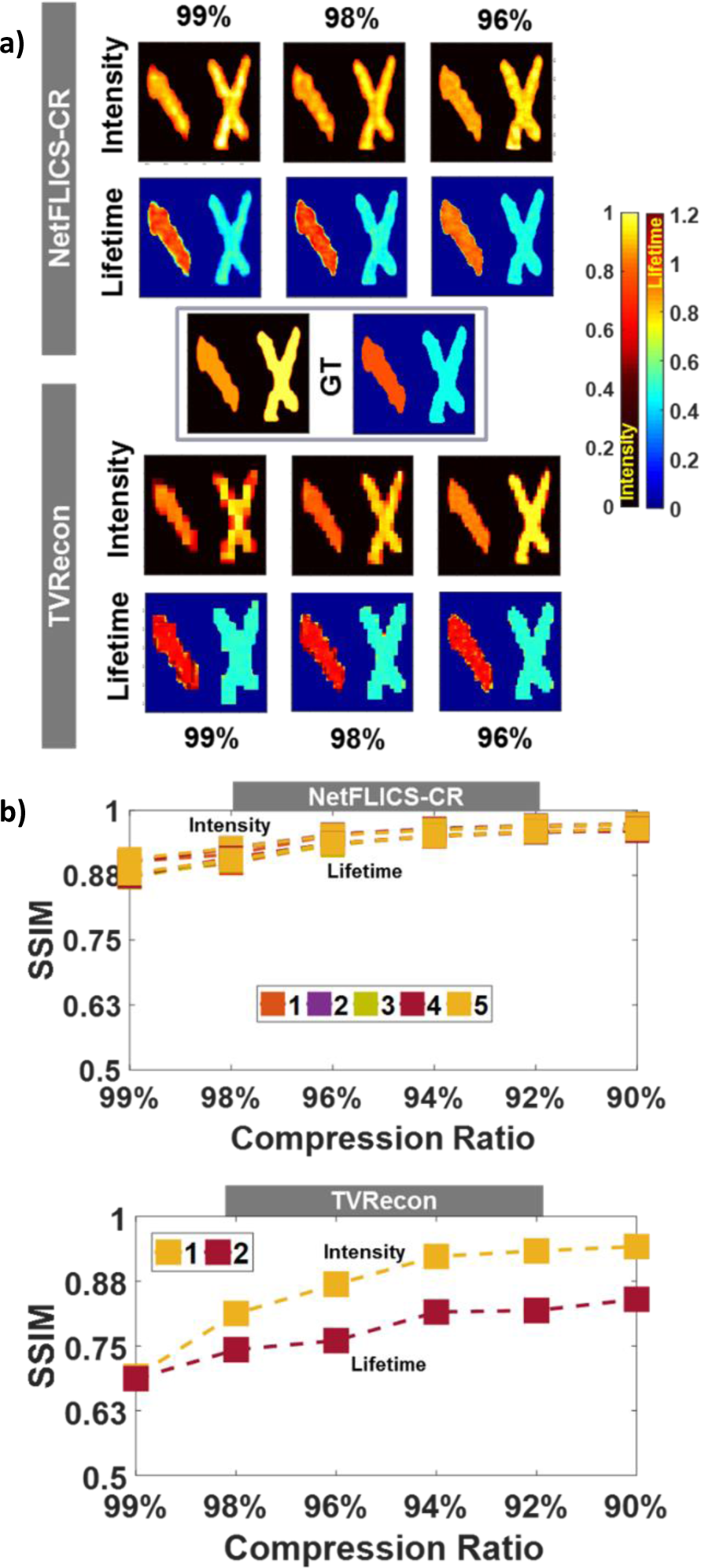
*In Silico* HMFLI Experiments: a) Intensity and lifetime reconstructions. One of 400 reconstructed samples displayed for visualization. b) Average intensity and lifetime SSIM values calculated versus ground-truth at different CRs: 90%-1,640, 92%-1,311, 94%-983, 96%-655, 98%-327 and 99%-163 out of 100%-16,384 patterns).

To test the experimental performance of NetFLICS-CR *in vitro* and the effect of compression on overall acquisition time, a sample composed of continuous letters “R”, “P” and “I” with respective volumes of 149, 124 and 81 µL was used. Alexa Fluor 750 (AF750; ThermoFisher Scientific, A33085) dye was prepared at 2.08 µM concentration to fill Letters R and I, while HITCI (Sigma Aldrich, 252034) at 40 µM was used for letter P. The latter should yield a higher lifetime value according to previous studies with this dye.^19,23^ The phantom was excited at 740 nm with 24 mW/cm^2^ power on a field of view of 35×35mm. The initial concentration of HITCI was monitored versus AF750 intensity under an external NIR CCD camera and was diluted with ethanol until matching the intensity of both dyes. The samples were acquired with the HMFLI system using an acquisition time of 0.5 s per pattern, for a total of ∼3 minutes for a 99% CR, ∼6 minutes for a 98% CR and ∼11 minutes for a 96% CR. Lower CRs were not considered as the main goal was to reduce experimental acquisition time through high data compression. Acquisition patterns were displayed on the digital micromirror device (DMD) as a positive and a negative part. The latter would be subtracted before reconstruction for noise removal. Therefore, respective pattern numbers of 163×2 for 99% CR, 327×2 for a 98% CR and 655×2 for a 96% CR were acquired. The raw data was reconstructed using both TVRecon and NetFLICS-CR workflows. For TVRecon reconstructions the above mentioned primary and secondary penalty parameters were used. For the lifetime quantification, TD pixels obtained from TVRecon with more than 5% counts of the maximum intensity value were fitted in order to obtain the lifetime values. Since this is a challenging minimization problem, the initial lifetime value was set to 0.8 ns and the fitting range bounded to [0.2-1.4] ns. This range covers the expected lifetime ranges of both letters at the same time. For NetFLICS-CR the *in silico* trained network was used at each CR to reconstruct both intensity and lifetime images (i.e. the network was not trained using experimental data). The reconstruction time of NetFLICS-CR was ∼2 s per spectral channel (16 spectral channels – total: ∼32 s), while the full TVRecon approach took ∼190 s per spectral channel (for a total of ∼50 minutes). After intensity reconstruction, for both methods, pixels with less than 20% of the maximum intensity value were set to 0 for background removal. Images were then normalized to the maximum value for SSIM comparison to an experimental intensity ground-truth (NIR CCD image of the sample plane).

The results for intensity and lifetime reconstructions for both methods are displayed in **Figure 3(a).** As shown in **Figure 3(b)**, SSIM values at 99%, 98% and 96% CRs are higher for NetFLICS-CR than for TVRecon reconstructions. In contrast to intensity, no lifetime ground-truth can be obtained, However, it is expected that due to the homogenous nature of the sample, the lifetime will be similar across a single letter. Hence, letters R and I should yield a similar lifetime as they contain the same fluorophore. Results for lifetime are displayed as histograms with a distribution per CR and method in **Figure 3(c)**. Two marked histogram peaks are located at average lifetimes of ∼1 ns and ∼0.5 ns, which respectively describe letters P (HITCI dye) and R/I (AF750 dye). For NetFLICS-CR, these average lifetimes are close to the expected values for the two fluorophores employed within this buffer.^19,23^ Conversely, at these CRs the TVRecon method returned a higher number of outliers than expected. Note that the pixels used for reporting the reconstructed lifetimes are the ones used for the intensity comparison. Hence, these *in vitro* results indicate that NetFLICS-CR, even if only model trained, retrieves more accurate lifetime and intensity values when applied to experimental data sets at high CRs. Additionally, NetFLICS-CR leads to a significant reduction in the acquisition time.

**Fig. 3.**
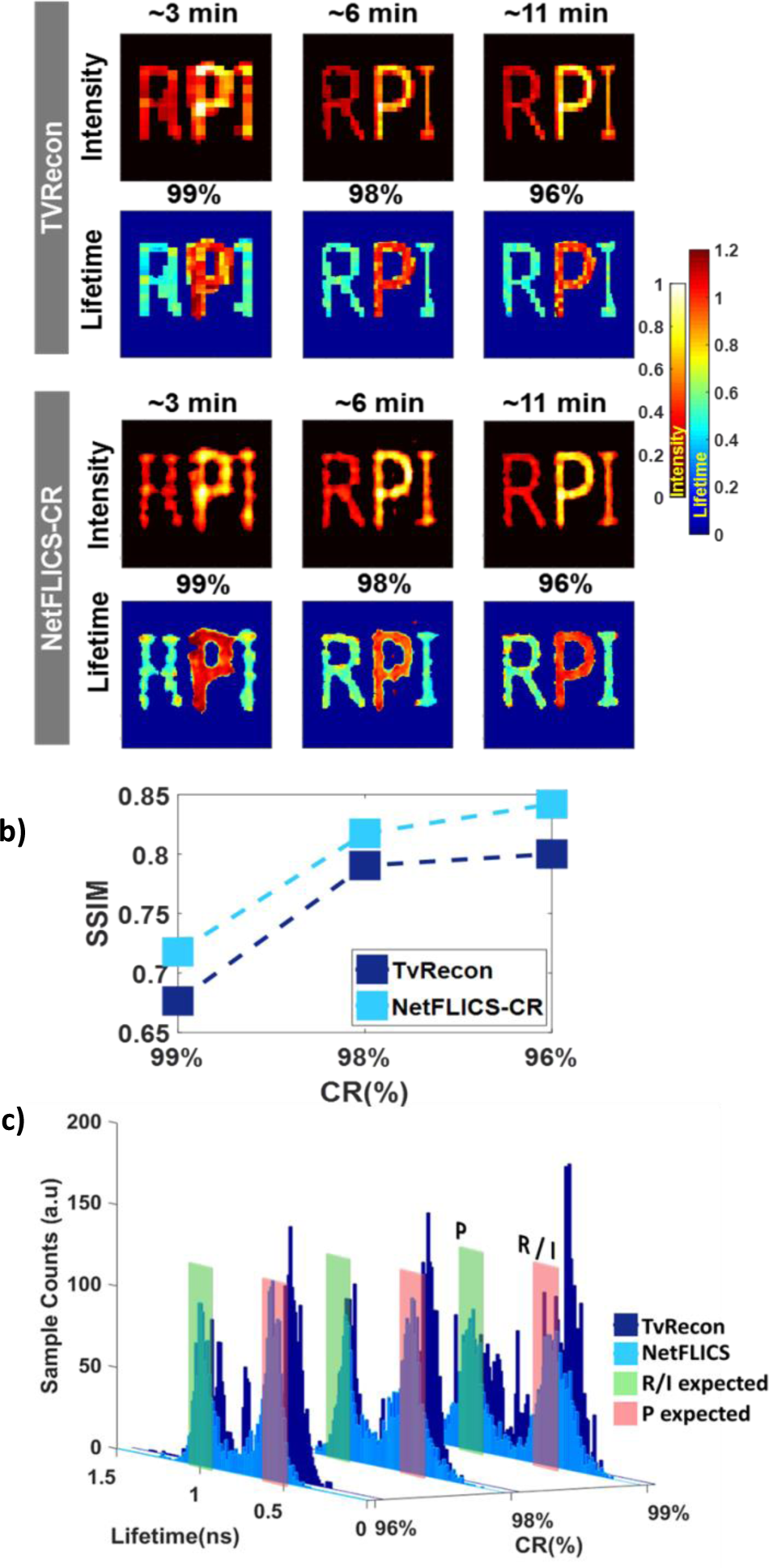
*In vitro* HMFLI Experiments: a) Intensity and lifetime reconstructions for RPI phantom per CR and acquisition time. b) SSIM values calculated comparing reconstructed versus external CCD ground-truth per CR. c) Distribution of lifetime reconstructions per CR and method for RPI phantom.

We further investigated the performances of NetFLICS-CR in a very challenging scenario, the quantification of receptor-ligand engagement and drug uptake in live, intact mice. This study was designed around two HMFLI imaging experiments that aim to: 1) Display lifetime differences between two organs through imaging of Transferrin (Tf) uptake in liver and bladder for a mouse injected with Tf-AF700 and imaged after 6 hours post-injection. 2) Reconstruct TZM binding to HER2-positive AU565 tumor xenograft upon intravenous injection of TZM-AF700 and TZM-AF750 via FRET imaging in two time-points. Sample preparation for both procedures is further explained in **Supplementary Material Section 3**. For the first procedure, the mouse was imaged at a minimum CR of 96% with ∼11 minutes acquisition time at 700 nm and 31 mW/cm^2^ excitation for a 38×38 mm FOV. Therefore, 99% and 98% CR reconstructions could be later retrieved to represent acquisitions of ∼6 and ∼3 minutes, respectively. Results for the Tf-AF700 mouse experiments are displayed in **Fig. S3**. Liver is a major site for iron homeostasis, so its transferrin receptor levels are high, leading to significant Tf uptake. In addition, due to the liver’s detoxifying function, changes in its microenvironment, such as in pH and ion composition, consistently result in a significant donor quenching even without the use of an acceptor probe (**Fig. S3**). In contrast, urinary bladder, an excretion organ, doesn’t cause donor fluorescent lifetime decrease, as it was shown by previous studies. ^28–30^ Both TVRecon and NetFLICS-CR methods were employed for comparison. NetFLICS-CR intensity and mean lifetime images were directly outputted from the network at CRs of 99%, 98% and 96%. On the other hand, TVRecon intensity images were reconstructed by TVAL3 inverse solver and mono-exponentially fitted through LSM to yield mean lifetime per pixel. Reconstructions have been overlaid over a grayscale intensity image of the FOV acquired with an external CCD camera. Intensity reconstructions have been normalized to maximum values in both methods for comparison. Intensity and lifetime reconstructions indicate that NetFLICS-CR better represents the expected biological behavior in liver and bladder, being capable of distinguishing the two organs by a change in lifetime. Especially, the bladder lifetime, which is an intra- and inter-subject control as FRET is not expected to occur in this organ due to the lack of receptor-mediated Tf uptake, is retrieved with good accuracy by NetFLICS-CR. Furthermore, it reports accurately on the lack of fluorescence expected at the 760nm control channel (Acceptor). Conversely, the TVRecon workflow fails in retrieving accurately both the intensity and lifetimes maps at these CRs and acquisition times.

To evaluate HMFLI + NetFLICS-CR ability to localize tumors and quantify *in vivo* drug uptake in dim conditions, AF700-TZM and AF750-TZM FRET pair was intravenously injected at a 2:1 acceptor to donor ratio in athymic nude mouse bearing an AU565 tumor xenograft that was imaged 24- and 102-hours post-injection. Further explanation of the xenograft preparation is shown in **Supplementary Material Section 3.** The mouse was imaged *in vivo* with a minimum CR of 98%. The excitation was set to 31 mW/cm^2^ at 700 nm for a 38×38 mm FOV, yielding a total acquisition time of ∼6 minutes (256 TD temporal bins and 16 spectral channels). Intensity and lifetime images are reconstructed per time point for both donor and acceptor peak channels at respective detection wavelengths of 719 nm and 760 nm. NetFLICS-CR intensity and mean lifetime were directly resolved, while TVRecon TD intensity data was inverse-solved by TVAL3. For TVRecon lifetime, each pixel was bi-exponentially fitted to return A1, A2, Tau1 and Tau2 values, which respectively represent the percentage of FRETing Donor (FD%), Acceptor and their lifetimes Tau1 and Tau2. To directly compare to the mean lifetime output of NetFLICS-CR, mean lifetime was calculated through **(*A1*/100)* *Tau1 +* (*A2*/100)* *Tau2.*** After reconstruction with both methods, intensity images were normalized and the region of interest defined by an external CCD intensity image of the sample plane. The resulting region of interest was also applied for lifetime reconstructions. FRET occurring between donor and acceptor fluorophores would lead to quenching of the donor intensity and reduction of donor lifetime by the acceptor at the targeted tumor area.^31^ Therefore, accessing both donor and acceptor channels could provide insights on the level and distribution of FRET events within the tumor.

NetFLICS-CR and TVRecon reconstructions at the tumor xenograft area per each time-point are shown in **Figure 4(a)** for 99% CR with their descriptive mean and standard deviation. A summary for the mean lifetime values at the tumor region per post-injection time, reconstruction method, detection channel and CR is summarized in **Figure 4(b)** and full set of reconstructed images displayed in **Fig. S4**. In order to control the descriptive statistics, means account for the highest 80-pixel values in the tumor area per method and reconstruction type. To the best of our knowledge, these are the first results reporting on the macroscopic assessment of *in vivo* intracellular delivery of TZM using HMFLI-based FRET imaging. ^32^According to the *in vitro* hyperspectral behavior of Tf-AF700 and Tf-AF750 and previously reported levels of FRET interaction between them, it is expected that the donor mean lifetime (expected value of ∼1 ns) will decrease as it is quenched by the acceptor (expected value of ∼0.5 ns), but the acceptor lifetime would minimally change.^16,19,33^

**Fig. 4.**
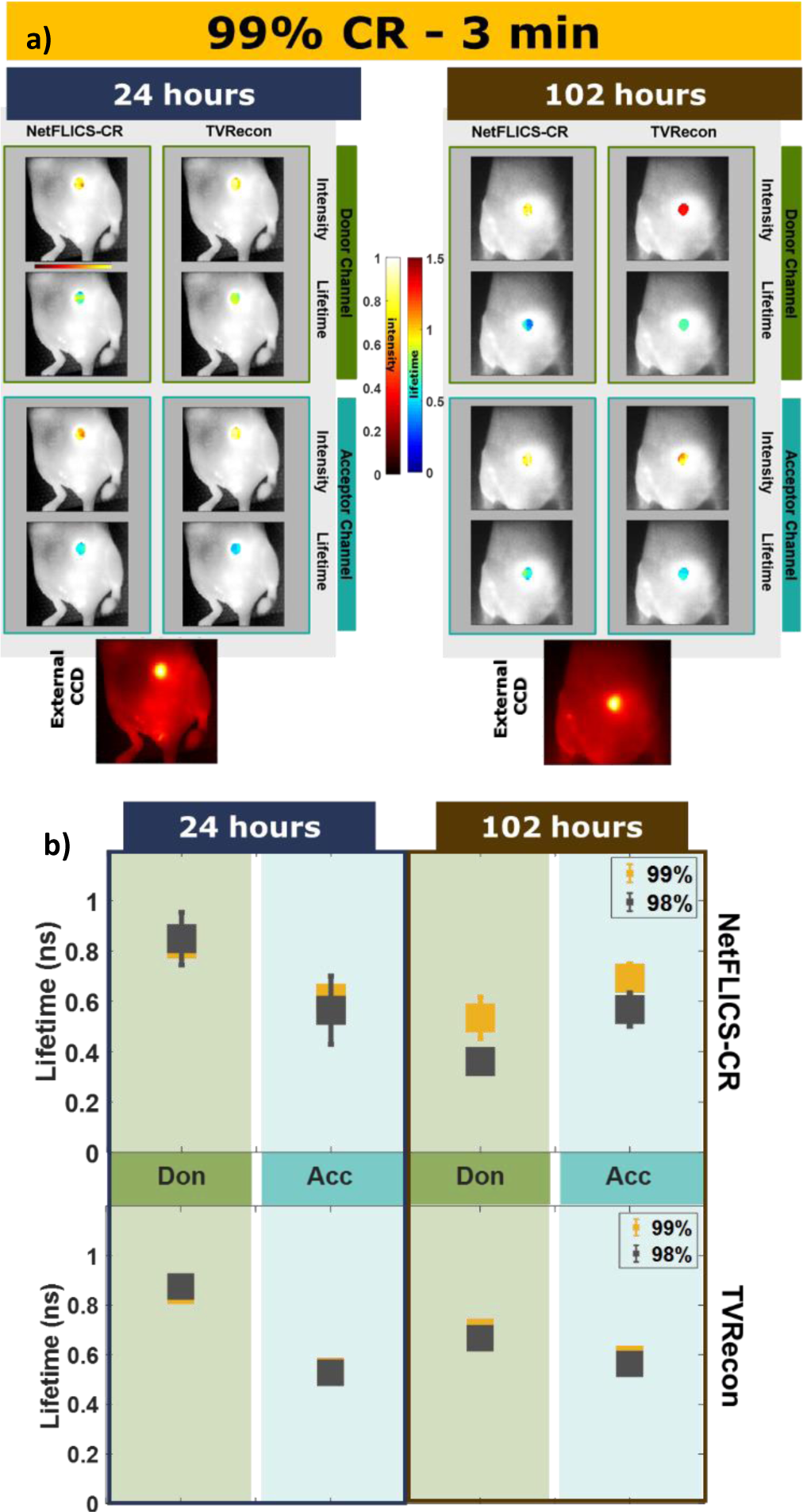
*In Vivo TZM experiments*: **a)** Intensity and lifetime reconstructions for the tumor region at 24- and 102-Hours post-injection at 99% CR. **b)** Mean lifetime and standard deviation for Donor and Acceptor channel per time point, reconstruction method and CR.

In order to validate the expected values for the peak wavelengths, individual AF700 donor and AF750 acceptor probes were quantified along detection channels as displayed in **Fig. S5** where lower detection channels are dominated by donor emission, while upper ones by acceptor emissions.As shown in **Figure 4(b)**, at 24 hours p.i. in the tumor region there is a decrease in lifetime from donor to acceptor channel. Considering ligand/target engagement in the xenograft region to be heterogenous, NetFLICS-CR better approximates the expected values at 99% and 98% CRs, where TVRecon shows regions with no variation, which could indicate an over-regularization despite optimal parameter selection. At 102 hours p.i. donor quenching is expected due to the binding of donor and acceptor labeled TZM to HER2 dimerized receptors, resulting in a decrease in donor lifetime, as the drug undergoes internalization and endocytic trafficking. Conversely, since lifetime is independent of intensity and concentration, even if the signal of the acceptor decreases, lifetime should minimally change. Of note, from 24 hrs to 102 hrs both the donor and acceptor channel raw fluorescence signals decrease as displayed in **Fig.S6**. This is expected as the fluorescently tagged drug is excreted from the live animals leading to reduced local concentrations overtime. Additionally, the decrease in donor lifetime from 24 hrs to 102 hrs and acceptor lifetime (**Fig.4(b)**) is suggestive of an increased fraction of AF700-TMZ undergoing FRET, i.e. intracellular delivery. This is expected since as the overall concentration of AF700-TMZ is decreasing due to excretion out of the animal, the intracellular fraction remaining is increasing over the extracellular fraction. Though, additional analysis and experiments of TZM engagement at the macroscopic level with immunohistochemistry validation are needed to have certainty of the amount of FRET expected at each timepoint. Last, even though Gaussian noise is included when simulating the fluorescent decays, we expect for future training sets to more accurately approximate the noise model and bi-exponential nature of the experimental TZM TPSFs; leading to overall improvements for lower photon count settings (tumor xenografts).

In conclusion, we report a novel CNN architecture, NetFLICS-CR, which efficiently reduces the acquisition time of high-dimensional HMFLI optical molecular data, while simultaneously producing 128×128 hyperspectral intensity and lifetime maps. Besides offering a fit free solution to the image formation paradigm, NetFLICS-CR led to a reduction in HMFLI acquisition times from ∼2.5 hours at 50% CR to ∼3 minutes at 99% CR. Despite the challenging photon-starved nature of *in vivo* acquisitions, NetFLICS-CR was able to reconstruct accurate intensity and lifetime maps that were in accordance with the expected biological outcome. Moreover, the image formation in NetFLICS-CR did not require any user input in contrast to fitting techniques, which paves the way to more standardized HMFLI for *in vivo* tissue characterization. Future work will pursue a further increase in resolution (beyond 128×128) as well as a decrease on the current ∼3-minute minimum acquisition time (at 0.5s exposure per pattern) through optimizing the system’s detection gating and binning parameters^34^. Beyond preclinical imaging, such a deep learning paradigm is expected to greatly facilitate the translation of these new analytical tools to the clinical settings where acquisition and processing times are critical for patient intervention, such as in optical guided surgery. Overall, this work illustrates how CNNs can play a central role in reducing the acquisition times for single-pixel multidimensional imaging at large, especially HMFLI. Additionally, we foresee that the NetFLICS-CR architecture herein proposed can be useful to guide other deep-learning developments in the field of compressed sensing and multiplexed imaging.

## Acknowledgments

We acknowledge the support of Jason T. Smith in improving lifetime accuracy. NIH grants R01EB19443/R01CA207725/R01CA237267. The NVIDIA Corporation for the donated Titan Xp GPU and the support of Genentech, Inc for donating the trastuzumab.

## Author Contributions

M.O., R.Y. and X.I. conceived the idea. M.O. and R.Y. designed the CNN and M.O. and X.I. the research study. M.O. performed the research study. A.R. performed the *in vivo* sample preparation. M.B. and X.I helped M.O. with biological interpretation of results. M.O. wrote the manuscript. X.I., A.R. and M.B. edited the manuscript. All authors discussed the conclusions.

