## Supplementary Material for "Deep Learning Enhanced Hyperspectral Fluorescence Lifetime Imaging"

Supplementary Information
for

### Deep Learning Enhanced Hyperspectral Fluorescence Lifetime Imaging

Marien Ochoa, Alena Rudkouskaya, Ruoyang Yao, Pingkun Yan, Margarida Barroso and Xavier Intes

 ,


#### 1. NetFLICS-CR Training and Architecture Specifics

Since the physics behind single-pixel measurement generation is known, NetFLICS-CR as well as NetFLICS are trained on simulated single-pixel acquisitions. EMNIST figures consisting of digits and letters are re-scaled from 28x28 pixels to the desired resolution, in the case of NetFLICS-CR, of 128x128 pixels. Furthermore, data is augmented by randomly clustering many of the rescaled EMNIST figures and then resizing the space to 128x128 by down-sampling. The images are rotated, flipped, rescaled and organized randomly across the 128x128 space so that no repeating figures exist. To mimic the single-pixel data generation, 128x128 intensity images randomly varying from 200 to 800 photon counts and their corresponding lifetime images with random values from 0.3 to 1.2 nanoseconds were used. These values aim to cover typical ranges obtained in the experimental sets. Since the time gates used in the experimental procedure are kept at 256 with intervals of 32.6 ps, a fluorescence decay curve with Poisson noise can be simulated for each pixel matching the given intensity and corresponding lifetime values of the “sample space”. TPSFs were “acquired” by convolving the decay with an experimentally acquired IRF. To simulate single-pixel acquisition, weights with range -1 to 1 given by the set of used “illumination Ranked Hadamard” patterns [1], are applied to the sum of all TPSFs from the “sample space”. As using the full pattern basis for a 128x128 resolution is not experimentally viable, a total of 1800 Ranked Hadamard patterns were used for data generation in order to cover a minimum CR of 90% for training. An example of these sets is shown in **Fig. S1**. A total of 40,000 sets were used, 32,000 for training and 8,000 for validation. After the samples are generated, the training per compression ratio is achieved by modifying the “pattern dimension” of the CW part of the data set as exemplified in **Fig. S1 (a)** for different CRs.

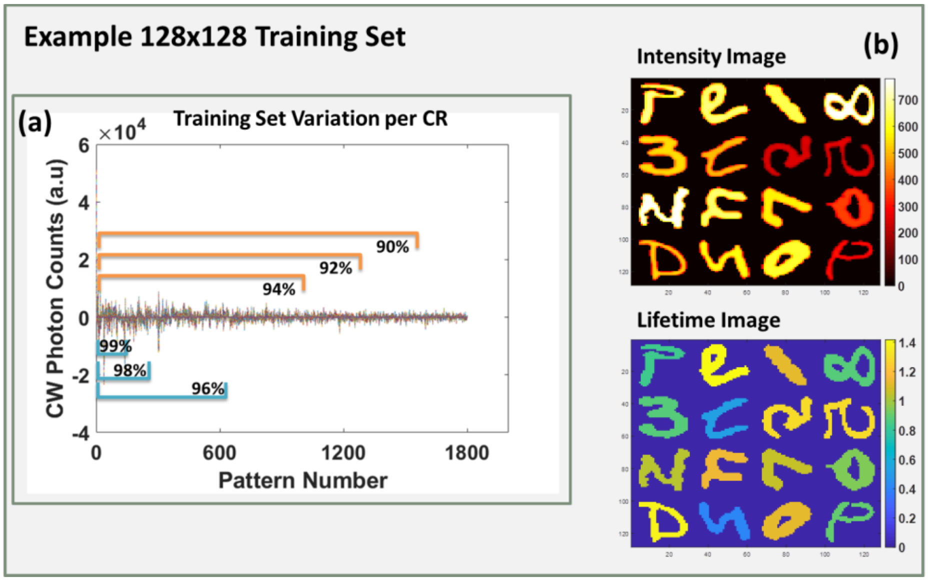


**Fig. S1.** (a) Continuous Wave visualization of simulated raw data and parts of the measurement that will be used to retrieve reconstructions for each compression ratio. (b) Example of simulated intensity and lifetime image corresponding to (a) for 128x128 resolution.

The architecture of NetFLICS-CR follows the same schematic as NetFLICS [2]. The common branch of NetFLICS-CR translates the input of size CNx256xPN, where PN represents the pattern number and CN number of detection channels, into 2D temporal data of dimensions 256 x 16384, which corresponds to the number of pixels in a TD 128x128 space. Additionally, in this branch, sparsity features are extracted from the input, through a 1D convolutional layer with 16384 size one 1D convolutional kernels that operate along the time dimension. Then batch normalization and ReLU activations are used. The output of segment 1 is permuted to 16384x256, so that it can be reshaped into the intensity branch to 128x128x256, which is one TPSF per pixel. One ResBlock of 256 kernels with size 3x3 is followed by a ReconBlock formed by respective kernel numbers and sizes of 64 and 1x1, 32 and 1x1 and 1 and 3x3, which results in the 128x128 intensity image per detection channel. Parallel to intensity reconstruction, the transposed output from the common segment is received by a 1D convolution with 512 kernels of size 1 and batch normalization/ReLU activation. The number of features increases from 256 to 512 in the same way as NetFLICS, which helps with the lifetime feature extraction. The output is further reshaped into 128x128x512, which is the input for a separable 2D convolution with 256 kernels of size 1x1 and followed by a ReLU activation[3]. Then a ResBlock and two ReconBlocks are used to yield the 128x128 lifetime image per detection channel. Compared to the original NetFLICS architecture, NetFLICS-CR takes longer to train, using the same GPU (NVIDIA Titan XP), due to the increase in number of kernels, a decrease in learning rate and an increase in the number of epochs.

#### 2. TVRecon Specifics

TVRecon is a method for intensity and lifetime image reconstruction composed of two main algorithms, TVAL3 inverse solver and Least-Squares Minimization (LSQR) algorithm. TVAL3 stands for "Total Variation Minimization by Augmented Langranian and Alternating Direction Algorithms", which is a Matlab inverse solver that helps retrieve original images from its degraded acquisitions. More information about TVAL3 mathematical model can be found in [4]. It is applicable to single-pixel imaging, where the sample plane is described by patterns with sparsity constraints. HMFLI input data is composed of a set of Time-Point-Spread-Function (TPSFs) and its recorded photon counts per detection wavelength over 256 time points. One of these sets is generated per recorded pattern. This data displayed in **Fig. S2 (a)** is the structure of the input for both TVRecon and NetFLICS-CR. Further explanation of the optical system and its arrangement to yield this type of output is provided in [5]. For TVRecon, the input is translated into one TPSF per pixel of the desired resolution by inverse solving ***m(t) = P*x(t)*** for ***x(t)*** as displayed in **Fig.S1 (b)**. The TVAL3 regularization terms used throughout this manuscript follow the isotropic model.

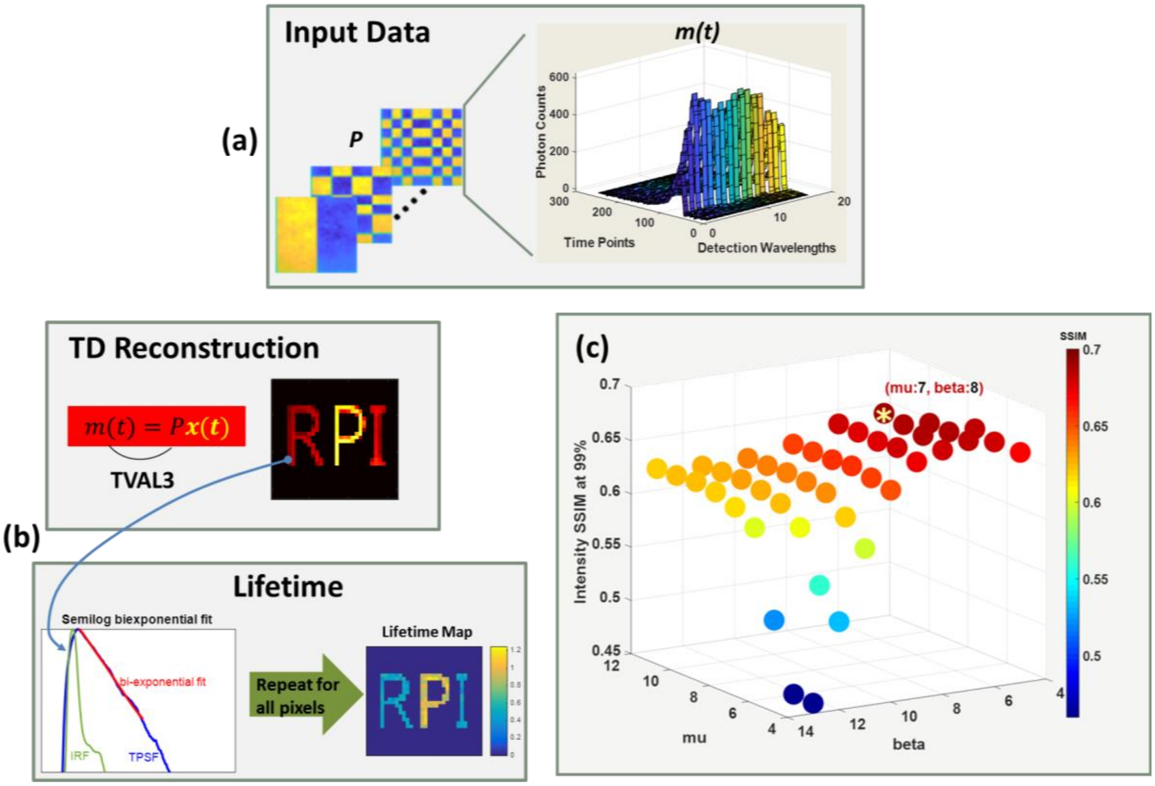


**Fig. S2.** (a) Input data for TVRecon and NetFLICS-CR methods. (b) Graphic description of TVRecon workflow. (c) Optimization of primary (mu) and secondary (beta) penalty parameters for TVAL3 based on intensity SSIM.

Since the algorithm is highly dependent on these terms, the primary and secondary penalty parameters, being the most important according to the developers [4], are tuned. According to the developers suggestion the primary penalty **μ** and secondary penalty **β** parameter should be ideally set between 2^4 and 2^13. Therefore, for the 400 samples used during simulation experiments Time-Domain reconstructions were made for all possible combinations of **μ** and **β** parameters. The Continuous-Wave intensity reconstructions were later evaluated versus ground-truth through SSIM. The combination yielding the highest SSIM value (closer to 1) was used for Time-Domain reconstructions. The SSIMs for the possible combinations are displayed in **Fig. S2 (c).** The estimated values were used for the other sets of experiments. For the second part of the TVRecon method, LSQR will be used to produce a fit and respective A1, A2, Tau1 and Tau2 values per each TPSF of the pixel space that will be further used in calculating the mean lifetime through **(A1/100)* Tau1 + (A2/100)* Tau2.** The optimization is done using Matlab's "fmincon" where upper and lower bounds have to be initialized. The initialization values correspond to the long and short lifetimes in nanoseconds, which are estimated factory values. Each value has a parameter bound with +/- units. These values are specified in the main text for each experimental set.

#### 3. In Vivo Experiments Sample Preparation

***Ligand labeling.*** Human holo Tf (Sigma) and Trastuzumab (Genentech) were conjugated to AF700 or AF750 (Life Technologies) through monoreactive N-hydroxysuccinimide ester to lysine residues in the presence of 100 mM Na bicarbonate, pH 8.3, according to manufacturer's instructions. The probes were purified by desalting columns and Amicon Ultra-4 microconcentrators (Millipore). The degree of labeling of the probes was assessed by spectrophotometer DU 640 (Beckman Coulter, Fullerton, CA, USA). The average degree of labeling was no more than 2 fluorophores per Tf or Trastuzumab molecule. All probes were normalized to concentration 1 mg/mL in phosphate-buffered saline pH 7.6 and filter sterilized.

*Cell culture*. Breast cancer cell line AU565 was purchased from ATCC (CRL-2351) and cultured in RPMI 1640 medium supplemented with 10% FBS in 5% $CO_2$ at 37$^oC$ in a humidified incubator for less than 12 passages.

*Animal experiments.* All animal procedures were conducted with the approval of the Institutional Animal Care and Use Committee at both Albany Medical College and Rensselaer Polytechnic Institute. Animal facilities of both institutions have been accredited by the American Association for Accreditation for Laboratory Animals Care International. Tumor xenografts were generated by injecting $10x10^6$ AU565 cells in phosphate-buffered saline (PBS) mixed 1:1 with Cultrex BME (R&D Systems Inc, Minneapolis, MN, USA) into the right inguinal mammary fat pad of female 4-week old athymic nude mice (NU(NCr)-Foxn1$^{nu}$, Taconic Biosciences, Rensselayer, NY, USA). The subcutaneous tumors were allowed to grow for 4-5 weeks and were monitored daily. Tf-AF700 (40 μg in the volume 100 μL) or TZM-AF700 and AF750 (20 μg and 40 μg respectively) were injected retro-orbitally in anesthetized animals. For imaging, mice were anesthetized using E-Z anesthesia breathing machine and placed on a heating pad to maintain body temperature. The depth of anesthesia was monitored by breathing rate and the absence of response to a foot pinch.

#### 4. In Vivo imaging Results

***Transferrin AF700.*** The Tf-AF700 *in vivo* results are displayed in **Fig. S3** for 99%, 98% and 96% CRs. The main objective is to distinguish between organs (liver or bladder) using lifetime reconstructions. Since the liver is a major organ for iron homeostasis, TfR levels in this detoxifying organ are very high. Due to the liver's detoxifying function, its microenvironment changes, such as in pH and ion composition, results in a significant donor quenching even without the use of an acceptor probe. In contrast, the urinary bladder, an excretion organ, doesn't cause donor fluorescent lifetime decrease. The Tf uptake in the liver results in a lifetime decrease compared to bladder, where the excreted probe accumulates. [6] The same trainings used for *in silico* and *in vitro* HMFLI experiments have been used for *in vivo* Tf-AF700 NetFLICS-CR reconstructions. For the TVRecon method, TVAL3 adopted the previously mentioned parameters and since only Tf-AF700 is used, lifetime was quantified through mono-exponential fitting with initial values of 0.9±0.5ns. After intensity and lifetime reconstructions for both methods, pixels with intensities lower than 50% of the maximum value were set to 0 for background removal purposes. As displayed in **Fig. S3** two detection channels were reconstructed. The "Donor" channel at 719nm should receive the highest emission from the AF700 probe, whereas for the "Acceptor" channel at 760nm little to no fluorescence should be reconstructed as there is no acceptor probe. This channel has been placed as a reconstruction control. Results indicate that NetFLICS-CR can better resolve the image sample plane by: (1) resolving the expected lifetime decrease in the liver compared to bladder from the original AF700 lifetime estimated in ~1ns as previously reported. (2) Resolving the low to none fluorescence lifetime levels expected at 760nm "Acceptor" detection. These tasks cannot be accomplished by the TVRecon method in the current optimization settings and CRs/acquisition times.

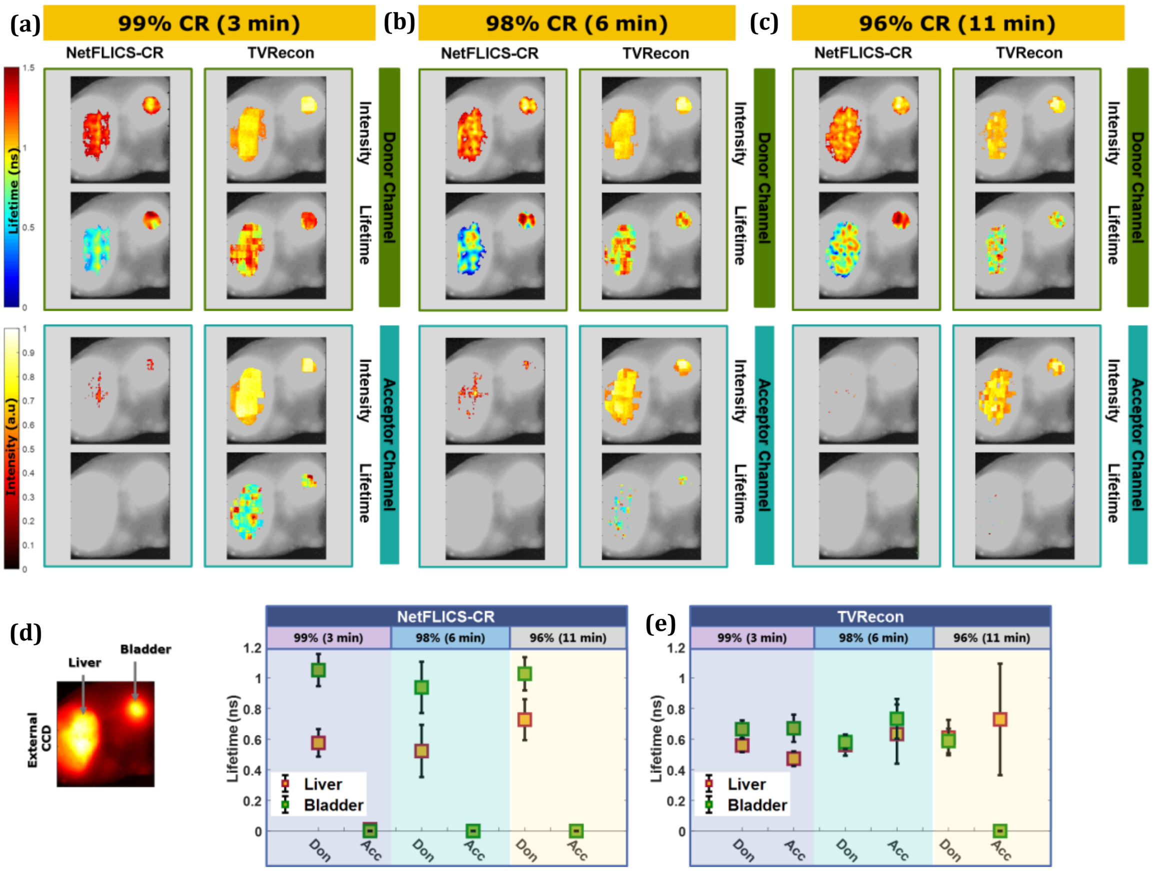


**Fig. S3.** Intensity and Lifetime reconstructions for Tf-AF700 Mouse experiments for 99% (a), 98% (b) and 96% (c) CRs for both NetFLICS-CR and TVRecon methods. Detection at 760nm for the Acceptor Channel is displayed as a control for reconstruction, since little detected fluorescence is expected at this wavelength. Liver and bladder areas are respectively indicated in (d), the external CCD image of the sample plane. Quantification for lifetime in liver and bladder areas per CR and reconstruction method (NetFLICS-CR or TVRecon) is summarized in (e) per CR, detection channel, organ type and reconstruction method.

***TZM AF700/AF750*.** Intensity and lifetime reconstructions for both 99% (3 minutes) and 98% (6 minutes) are displayed in **Fig.S4** for both methods and peak donor/acceptor channels.

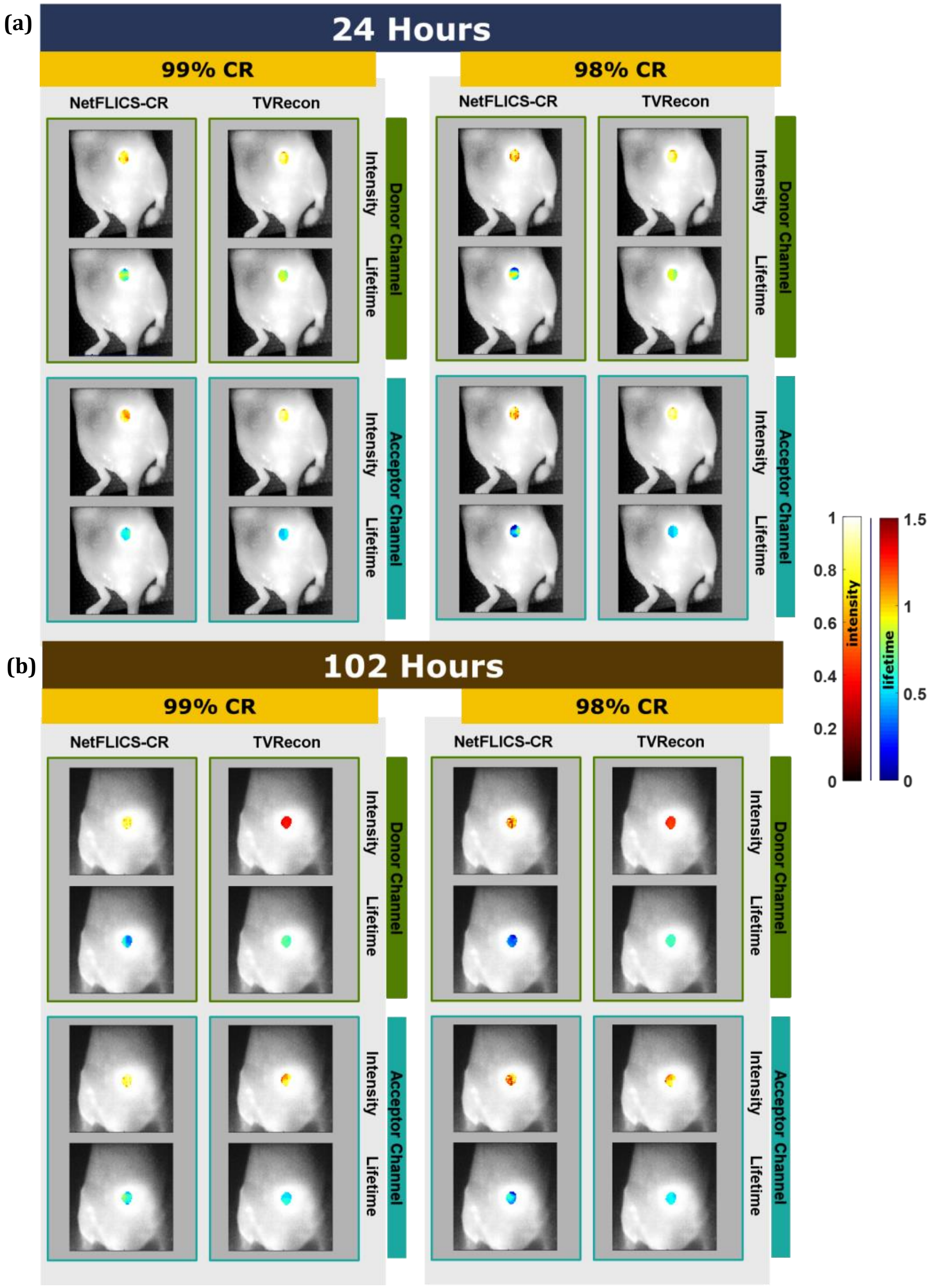


**Fig. S4 | *In Vivo TZM experiments reconstructions*:** Intensity and lifetime reconstructions for an athymic nude mouse with TZM targeted tumor xenograft at 24 (a) and 102 (b) hours post-injection. Reconstructions provided for both Donor (green) and Acceptor (Teal) channels for a 99% and 98% CR.

Further *in vivo* experiments with TZM FRET need to be done to fully understand its physics at the macroscopic level. For the purpose of this paper and considering lifetime values might change during probe preparation due to conjugation and the buffer solution, we have measured *in vitro* AF700 and AF750 fluorophores across 16 detection channels, as shown in **Fig.S5.** For lifetime, **Fig.S5 (b)** shows the mean and standard deviation for the peak Donor (0.90±0.10ns) and Acceptor (0.43±0.20ns) detection channels, which are subsequently used as the expected values for the *in vivo* fluorescence experiments. Concentrations of AF700 and AF750 followed the previously described 20 μg and 40 μg concentrations used to yield a 2:1 ratio. Both average intensities and lifetimes are displayed per each detection channel ranging from channel 1 at 715nm to channel 16th at 783nm, with 4.5nm wavelength space between channels.

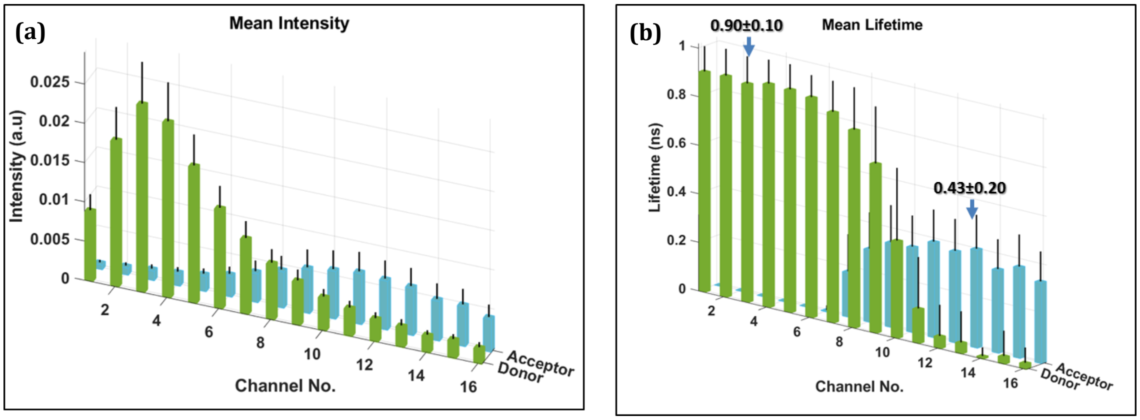


**Fig. S5.** The average intensity values for both donor and acceptor are displayed in Figure (a). Mean lifetimes and their standard deviations are shown per detection channel in (b). Lifetime values in peak channels used for reconstruction are emphasized.

To further understand the effect of post-injection time on the targeted fluorescent xenograft region, the raw TCSPC acquired data is analyzed by peak spectral donor and acceptor channel. Since the Hadamard Ranked basis is organized by spatial frequency, Pattern 1 is expected to yield the signal with the most intense fluorescence decay. The TPSFs are plotted per donor and acceptor channel at the two different time points as displayed in **Fig. S6.** At 24 hours the TPSF at the donor channel is less intense than the TPSF at the acceptor channel. On the contrary, at 102 hours post injection, despite using the same acquisition settings used at 24 hours (excitation at 700 nm and 0.031 W/cm$^2$ for a 38x38 mm FOV), both donor and acceptor TPSFs are below 200 photon counts. Despite the decrease in counts, both methods yielded respective reconstructions as shown in **Fig. S4.** Future research is needed to understand which is the lower photon count threshold that can be reconstructed for both methods and how it varies per experimental setting.

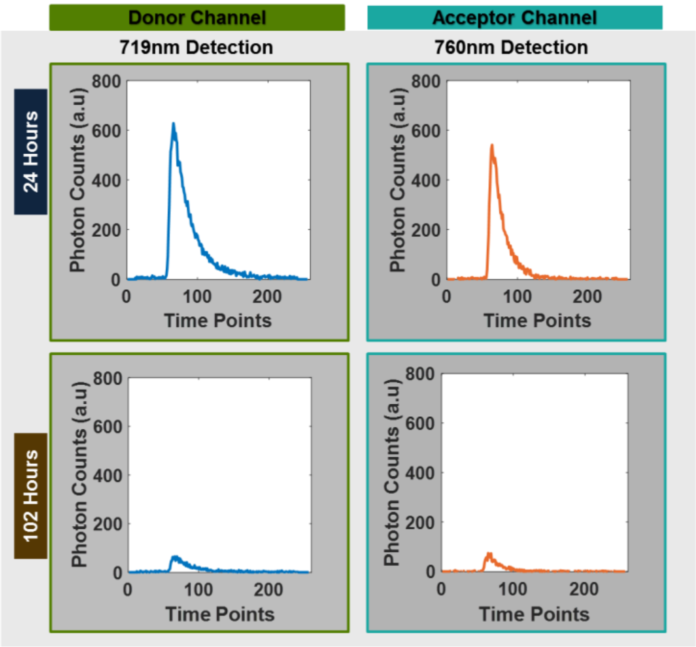


**Fig. S6.** Raw TPSF for Pattern 1 for both Donor and Acceptor channels at respective wavelengths of ~719nm and ~760nm. Since the Hadamard Ranked basis is organized by spatial frequency, Pattern 1 is expected to yield the signal with the highest fluorescence intensity. Pattern 1 TPSF is displayed per channel for 24 and 102 Hours raw TZM in vivo acquisitions.

[1] M. Ochoa, Q. Pian, R. Yao, N. Ducros, and X. Intes, “Assessing patterns for compressive fluorescence lifetime imaging,” *Opt. Lett.*, vol. 43, no. 18, pp. 4370–4373, 2018.

[2] R. Yao, M. Ochoa, P. Yan, and X. Intes, “Net-FLICS: fast quantitative wide-field fluorescence lifetime imaging with compressed sensing–a deep learning approach,” *Light Sci. Appl.*, vol. 8, no. 1, p. 26, 2019.

[3] J. T. Smith *et al.*, “Ultra-fast fit-free analysis of complex fluorescence lifetime imaging via deep Learning,” *bioRxiv*, p. http://dx.doi.org/10.1101/523928, 2019.

[4] C. Li, “Compressive sensing for 3D data processing tasks: applications, models and algorithms,” 2011.

[5] Q. Pian, R. Yao, N. Sinsuebphon, and X. Intes, “Compressive hyperspectral time-resolved wide-field fluorescence lifetime imaging,” *Nat. Photonics*, vol. 11, no. 7, p. 411, 2017.

[6] K. Abe, L. Zhao, A. Periasamy, X. Intes, and M. Barroso, “Non-invasive in vivo imaging of near infrared-labeled transferrin in breast cancer cells and tumors using fluorescence lifetime FRET,” *PLoS One*, vol. 8, no. 11, p. e80269, 2013.
